# Septate junctions regulate gut homeostasis through regulation of stem cell proliferation and enterocyte behavior in *Drosophila*

**DOI:** 10.1101/582148

**Authors:** Yasushi Izumi, Kyoko Furuse, Mikio Furuse

## Abstract

Smooth septate junctions (sSJs) contribute to the epithelial barrier, which restricts leakage of solutes through the paracellular route of epithelial cells in the *Drosophila* midgut. We previously identified three sSJ-associated membrane proteins, Ssk, Mesh, and Tsp2A, and showed that these proteins were required for sSJ formation and intestinal barrier function in the larval midgut. Here, we investigated the roles of sSJs in the *Drosophila* adult midgut. Depletion of any of the sSJ-proteins from enterocytes resulted in remarkably shortened lifespan and intestinal barrier dysfunction in flies. Interestingly, the sSJ protein-deficient flies showed intestinal hypertrophy accompanied by accumulation of morphologically abnormal enterocytes. The phenotype was associated with increased stem cell proliferation and activation of the MAP kinase and Jak-Stat pathways in stem cells. Loss of cytokines Unpaired2 and Unpaired3, which are involved in Jak-Stat pathway activation, suppressed the intestinal hypertrophy, but not the increased stem cell proliferation, in flies lacking Mesh. The present findings suggest that SJs play a crucial role in maintaining tissue homeostasis through regulation of stem cell proliferation and enterocyte behavior in the *Drosophila* adult midgut.

**Summary statement:** Depletion of smooth septate junction-associated proteins from enterocytes in the *Drosophila* adult midgut results in intestinal hypertrophy accompanied by accumulation of morphologically aberrant enterocytes and increased stem cell proliferation.

## Introduction

The intestinal epithelium serves as a physical barrier that prevents infiltration of food-derived harmful substances, microbial contaminants, and digestive enzymes into the body. To constitute an effective intestinal barrier, specialized cell-cell junctions, namely occluding junctions, play a crucial role in regulating free diffusion of solutes through the paracellular route. Septate junctions (SJs) are occluding junctions in invertebrates and act as the functional counterparts of vertebrate tight junctions (Anderson and Van Itallie, 2009, Furuse and Tsukita, 2006, Lane, 1994, Tepass and Hartenstein, 1994). In arthropods, morphological variation of SJs has been observed among different types of epithelia (Lane, 1994, Tepass and Hartenstein, 1994). Most ectodermally-derived epithelia and the perineural sheath have pleated SJs (pSJs), while endodermally-derived epithelia, including the midgut, have smooth SJs (sSJs) (Lane, 1994, Tepass and Hartenstein, 1994). Genetic and molecular analyses in *Drosophila* have revealed a number of molecular components and functional properties of pSJs (Tepass et al., 2001, Wu and Beitel, 2004, Banerjee et al., 2006, Izumi and Furuse, 2014). In contrast, few studies have been carried out on sSJs, and their physiological role is not well understood. Recently, we identified three sSJ-specific membrane proteins: Ssk, Mesh, and Tsp2A. Ssk is a four transmembrane domain-containing protein (Yanagihashi et al., 2012). Mesh is a single-pass membrane protein containing a large extracellular region (Izumi et al., 2012). Tsp2A belongs to the tetraspanin family (Izumi et al., 2016). Zygotic loss of any of these proteins results in embryonic lethality just before hatching or at the 1st instar larva stage with impairment of sSJ formation as well as epithelial barrier function, suggesting critical roles of sSJs (Yanagihashi et al., 2012, Izumi et al., 2012, Izumi et al., 2016, Izumi and Furuse, 2014, Furuse and Izumi, 2017).

The *Drosophila* adult midgut epithelium is composed of absorptive enterocytes (ECs), secretory enteroendocrine cells (EEs), intestinal stem cells (ISCs), EC progenitors (enteroblasts: EBs), and EE progenitors (enteroendocrine mother cells: EMCs) (Micchelli and Perrimon, 2006, Ohlstein and Spradling, 2006, Guo and Ohlstein, 2015). The sSJs are formed between adjacent ECs and between ECs and EEs (Resnik-Docampo et al., 2017). ECs and EEs are continuously renewed by proliferation and differentiation of the ISC lineage through the production of intermediate differentiating cells, EBs and EMCs. This renewal of adult midgut epithelial cells is critical for maintenance of homeostasis in the adult midgut. Recent studies have suggested that sSJs influence proliferation and differentiation of the ISC lineage in the adult midgut. Experimental suppression of Gliotactin, a tricellular junction-associated protein, in ECs led to epithelial barrier dysfunction, increased ISC proliferation, and blockade of EC differentiation in young flies (Resnik-Docampo et al., 2017). Moreover, loss of *mesh* and *Tsp2A* in clones caused defects in polarization and integration of ECs in the adult midgut (Chen et al., 2018). Thus, it will be interesting to examine the role of sSJs in the adult midgut in terms of the regulation of ISC proliferation and tissue homeostasis by genetic ablation of the sSJ-protein genes throughout ECs.

Here, we describe that depletion of the sSJ-proteins Ssk, Mesh, and Tsp2A from ECs causes remarkably reduced lifespan and midgut barrier dysfunction in flies. The sSJ protein-deficient flies show intestinal hypertrophy accompanied by accumulation of morphologically aberrant ECs and increased stem cell proliferation. Interestingly, we show that the interleukin-6-like cytokines Unpaired2 and/or Unpaired3 are involved in this intestinal hypertrophy. We conclude that sSJs are crucial for the regulation of stem cell proliferation and EC behavior in the *Drosophila* adult midgut.

## Results

### Depletion of sSJ-proteins from ECs in adult flies results in shortened lifespan and midgut barrier dysfunction

To investigate the effect of sSJ-protein depletion on the *Drosophila* adult midgut, UAS-RNAi lines for sSJ-proteins were expressed in ECs using a *Myo1A*-Gal4 driver, combined with a temporal and regional gene expression targeting (TARGET) system (Zeng et al., 2010). The UAS-RNAi lines used for the sSJ-proteins were UAS-*ssk*-RNAi, *mesh*-RNAi (15074R-1), and *Tsp2A*-RNAi (11415R-2), which were confirmed to effectively reduce the expression of their respective sSJ-proteins (Yanagihashi et al., 2012, Izumi et al., 2012, Izumi et al., 2016). *Myo1A*-Gal4 tubGal80^ts^ UAS-*CD8-GFP* and UAS-*ssk*-RNAi, *mesh*-RNAi, or *Tsp2A*-RNAi (*Myo1A^ts^*> *ssk*-RNAi, *mesh*-RNAi, or *Tsp2A*-RNAi) flies were raised to adults at 18°C and then shifted to 29°C to express each UAS-driven transgene. Almost all flies expressing the RNAis targeting the sSJ-protein transcripts (hereafter referred to as sSJp-RNAis) died within 10 days after transgene induction, while more than 95% of control flies (*Myo1A^ts^*> *CD8-GFP*) survived until 15 days after induction (Fig. 1A). Thus, reduced expression of sSJ-proteins in ECs results in remarkably shortened lifespan in adult flies. Next, we examined whether the barrier function of the midgut was disrupted in sSJp-RNAis flies. According to the method of a barrier integrity assay/Smurf assay method (Rera et al., 2011, Rera et al., 2012), flies were fed a non-absorbable 800-Da blue food dye in sucrose solution and observed for leakage of the dye from the midgut. At 3 days after transgene induction, reduced expression of any of the sSJ-proteins in ECs led to a significant increase in flies with blue dye throughout their body cavity, indicating defective midgut barrier function (Fig. 1C). Flies with midgut barrier dysfunction were further increased at 5 days after transgene induction, compared with age-matched controls (Fig. 1B, C). Thus, we confirmed that sSJ-proteins are required for the barrier function in the adult midgut, similar to the observations in the larval midgut (Yanagihashi et al., 2012, Izumi et al., 2012, Izumi et al., 2016).

**Figure 1.**
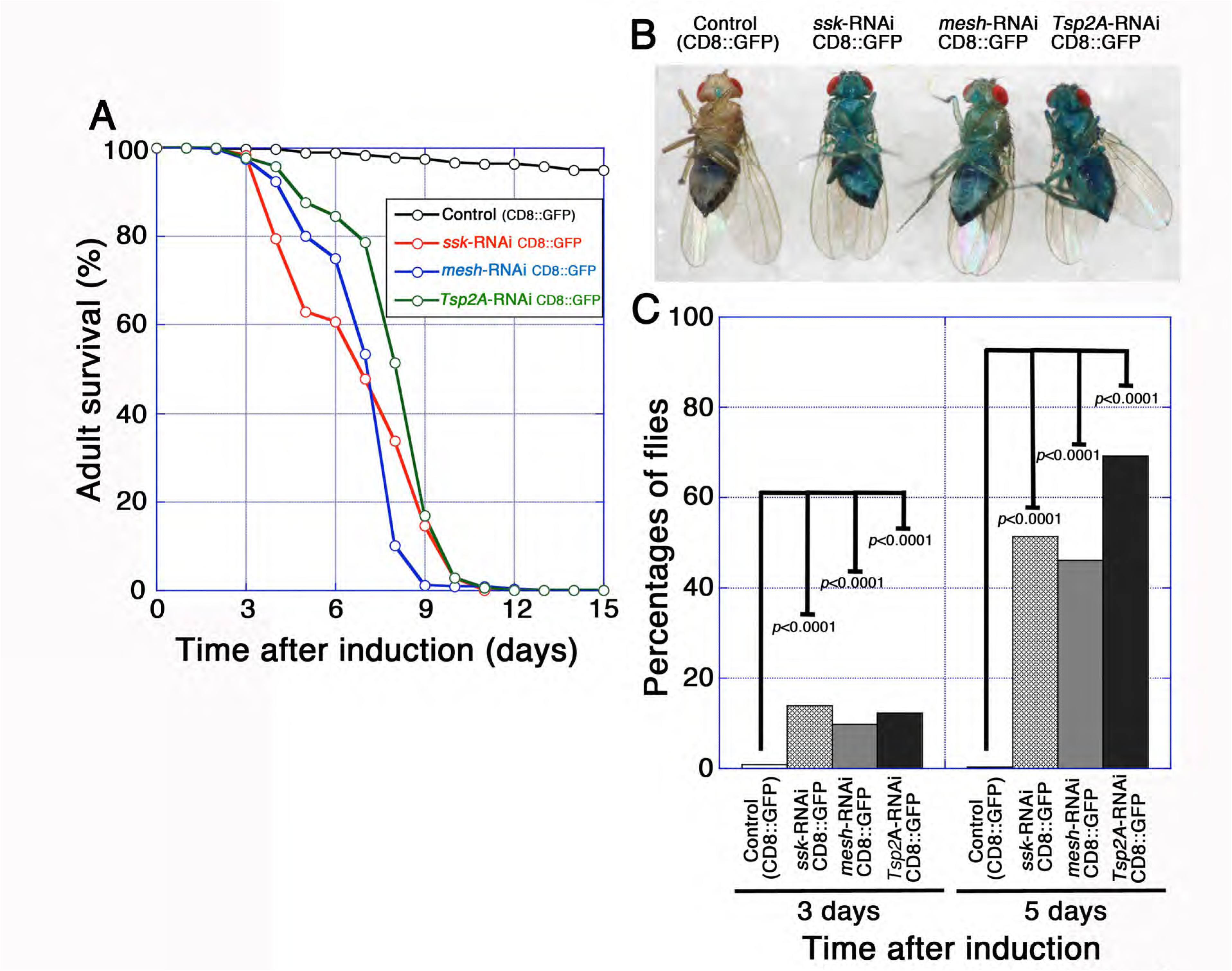
Depletion of sSJ-proteins from ECs in adult flies results in shortened lifespan and midgut barrier dysfunction. **(A)** Survival analysis of flies expressing *Myo1A^ts^*-Gal4/UAS-*CD8-GFP* without (control, *n*=240) or with UAS-*ssk*-RNAi (*n*=220), UAS-*mesh*-RNAi (15074R-1) (*n*=239), or UAS-*Tsp2A*-RNAi (11415R-2) (*n*=220). The transgenes were expressed with temperature-sensitive GAL80, and thus the flies were raised at 18°C until adulthood and then moved to 29°C. Each vial contained 20 flies (10 females, 10 males). **(B, C)** Barrier integrity assays (Smurf assays). Flies expressing *Myo1A^ts^*-Gal4/UAS-*CD8-GFP* without or with UAS-*ssk*-RNAi, UAS-*mesh*-RNAi, or UAS-*Tsp2A*-RNAi were fed blue dye in sucrose solution. (B) Typical examples of the phenotypes at 5 days after transgene induction. (C) Left to right: Control (*CD8-GFP*) (*n*=264), *ssk*-RNAi *CD8-GFP* (*n*=375), *mesh*-RNAi *CD8-GFP* (*n*=531), and *Tsp2A*-RNAi *CD8-GFP* (*n*=508) at 3 days after induction, Control (*CD8-GFP*) (*n*=232), *ssk*-RNAi *CD8-GFP* (*n*=299), *mesh*-RNAi *CD8-GFP* (*n*=384), and *Tsp2A*-RNAi *CD8-GFP* (*n*=336) at 5 days after induction. Loss of midgut barrier function was determined when dye was observed outside the midgut. Flies with reduced sSJ-protein expression show loss of barrier function compared with control flies (*CD8-GFP* flies). The *p*-values in (C) represent significant differences in pairwise post-test comparisons indicated by the corresponding bars (Fisher’s exact test).

### Depletion of sSJ-proteins from ECs leads to intestinal hypertrophy accompanied by accumulation of morphologically aberrant ECs in the midgut

The shortened lifespan and midgut barrier dysfunction in sSJ-protein-deficient flies prompted us to examine the organization of their midgut epithelium. At 5 days after transgene induction, a typical simple epithelium in which ECs expressed CD8-GFP driven by *Myo1A*-Gal4 was observed in the control midgut (Fig. 2A, E). Intriguingly, the organization of the epithelium was severely disrupted in the sSJp-RNAis midgut: a large number of ECs failed to become integrated into the epithelium and instead accumulated throughout the midgut lumen (Fig. 2B–D). In addition, the ECs exhibited a variety of aberrant appearances, implying a polarity defect (Fig. 2B–D). Indeed, abnormal distributions of actin and Dlg, a cell polarity protein, were observed in sSJp-RNAis ECs (Fig. 2B–D and Fig. S1). The most posterior part of the midgut had a hypertrophic phenotype: the lumen was filled with a large number of morphologically aberrant ECs and the diameter was severely expanded (Fig. 2F–I). We confirmed that expression of sSJp-RNAis in ECs led to decreased levels of the respective target proteins and mislocalization of other sSJ-proteins in the midgut (Fig. S2). Additional RNAi lines for *mesh* and *Tsp2A* showed essentially the same phenotypes (Fig. S3). Toluidine blue staining of semi-thin sections and ultrastructural analysis by transmission electron microscopy confirmed that morphologically aberrant cells were stratified in the sSJp-RNAis midgut (Fig. 2K–M, O–Q), while a monolayer of ECs with developed microvilli facing the lumen was observed in the control midgut (Fig. 2J, N). Notably, microvilli-like structures were often observed between stratified ECs in the sSJp-RNAis midgut (Fig. 2O, Q–S). Thus, depletion of any of the sSJ-proteins from ECs causes intestinal hypertrophy accompanied by dysplasia-like accumulation of ECs in the midgut lumen, suggesting that sSJs are required for homeostasis of the midgut organization.

**Figure 2.**
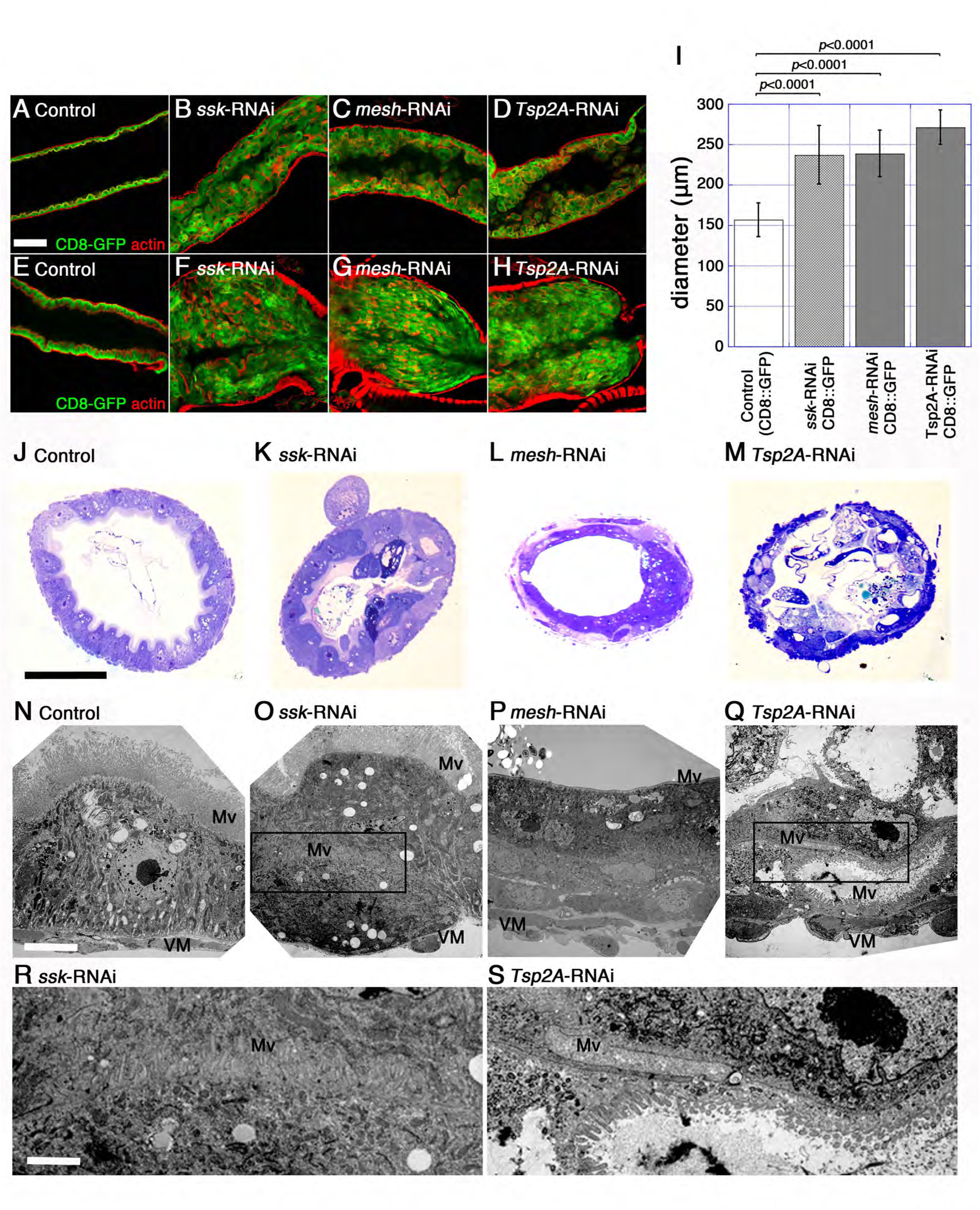
Depletion of sSJ-proteins from ECs leads to intestinal hypertrophy accompanied by accumulation of morphologically aberrant ECs in the midgut. **(A–H)** Confocal images of the adult midgut expressing *Myo1A^ts^*-Gal4/UAS-*CD8-GFP* without (A, E, control) or with UAS-*ssk*-RNAi (B, F), UAS-*mesh*-RNAi (C, G), or UAS-*Tsp2A*-RNAi (D, H) at 5 days after induction stained for actin (red). CD8-GFP driven by *Myo1A^ts^* was expressed in the ECs of each midgut. A large number of ECs with aberrant morphology have accumulated in the anterior (B–D) and posterior (F–H) midgut. The most posterior region of the sSJp-RNAis midgut is severely expanded (E–H). Scale bar: 50 μm. **(I)** Diameter of the most posterior region of the midgut. The diameter of the midgut was measured just anterior to the Malpighian tubules. Left to right: Control (*CD8-GFP*) (*n*=19), *ssk*-RNAi *CD8-GFP* (*n*=24), *mesh*-RNAi *CD8-GFP* (*n*=19), and *Tsp2A*-RNAi *CD8-GFP* (*n*=22) at 5 days after induction. Error bars show s.e.m. Statistical significance (*p*<0.0001) was evaluated by Student’s *t*-test. (**J–M**) Toluidine blue staining of the adult female anterior midgut in control (*CD8-GFP*) (J), *ssk*-RNAi *CD8-GFP* (K), *mesh*-RNAi *CD8-GFP* (L), and *Tsp2A*-RNAi *CD8-GFP* (M) flies at 5 days after induction. Stratification of cells in the midgut lumen is observed in the sSJp-RNAis midgut. **(N–S)** Transmission electron microscopy of the adult female anterior midgut in control (*CD8-GFP*) (N), *ssk*-RNAi *CD8-GFP* (O, R), *mesh*-RNAi *CD8-GFP* (P), and *Tsp2A*-RNAi *CD8-GFP* (Q, S) flies at 5 days after induction. (R) and (S) are enlarged views of the regions outlined by the black boxes in (O) and (Q), respectively. Morphologically aberrant cells are stratified in the sSJp-RNAis midgut (K–M). Microvilli-like structures are found between stratified ECs in the sSJp-RNAis midgut (O, Q–S). Scale bars: 5 μm (N–Q); 15 μm (R, S). Mv, microvilli; VM, visceral muscles.

### Depletion of sSJ-proteins from ECs leads to increased ISC proliferation in the midgut

We speculated that the EC accumulation was caused by overproduction of ECs in the sSJp-RNAis midgut. Because regeneration of ECs depends on proliferation and differentiation of the ISC linage, we examined whether proliferation of ISCs was increased in the sSJp-RNAis midgut. We stained the midgut with an antibody against phospho-histone H3 (PH3), a mitotic marker, and found that PH3-positive cells were markedly increased in the sSJp-RNAis midgut compared with the control midgut (Fig. 3A–D, M). Furthermore, immunostaining of the midgut with an antibody against Delta, an ISC marker, revealed that ISCs were increased in the sSJp-RNAis midgut compared with the control midgut (Fig. 3A–D, M). We also confirmed that PH3-positive cells were Delta-positive (Fig. 3A–D). Furthermore, the number of cells expressing Escargot (Esg)-LacZ, an ISC/EB marker, was increased in the *ssk-* or *mesh*-RNAi midgut (Fig. S4A–C). These results indicate that reduced expression of sSJ-proteins in ECs leads to increased ISC proliferation. Of note, Esg-LacZ signals were often observed in cells expressing CD8-GFP driven by *Myo1A*-GAL4 in the *ssk-* or *mesh*-RNAi midgut (Fig. S4E–F’’), suggesting that sSJ-protein depletion causes mis-differentiation of the ISC linage in the midgut.

**Figure 3.**
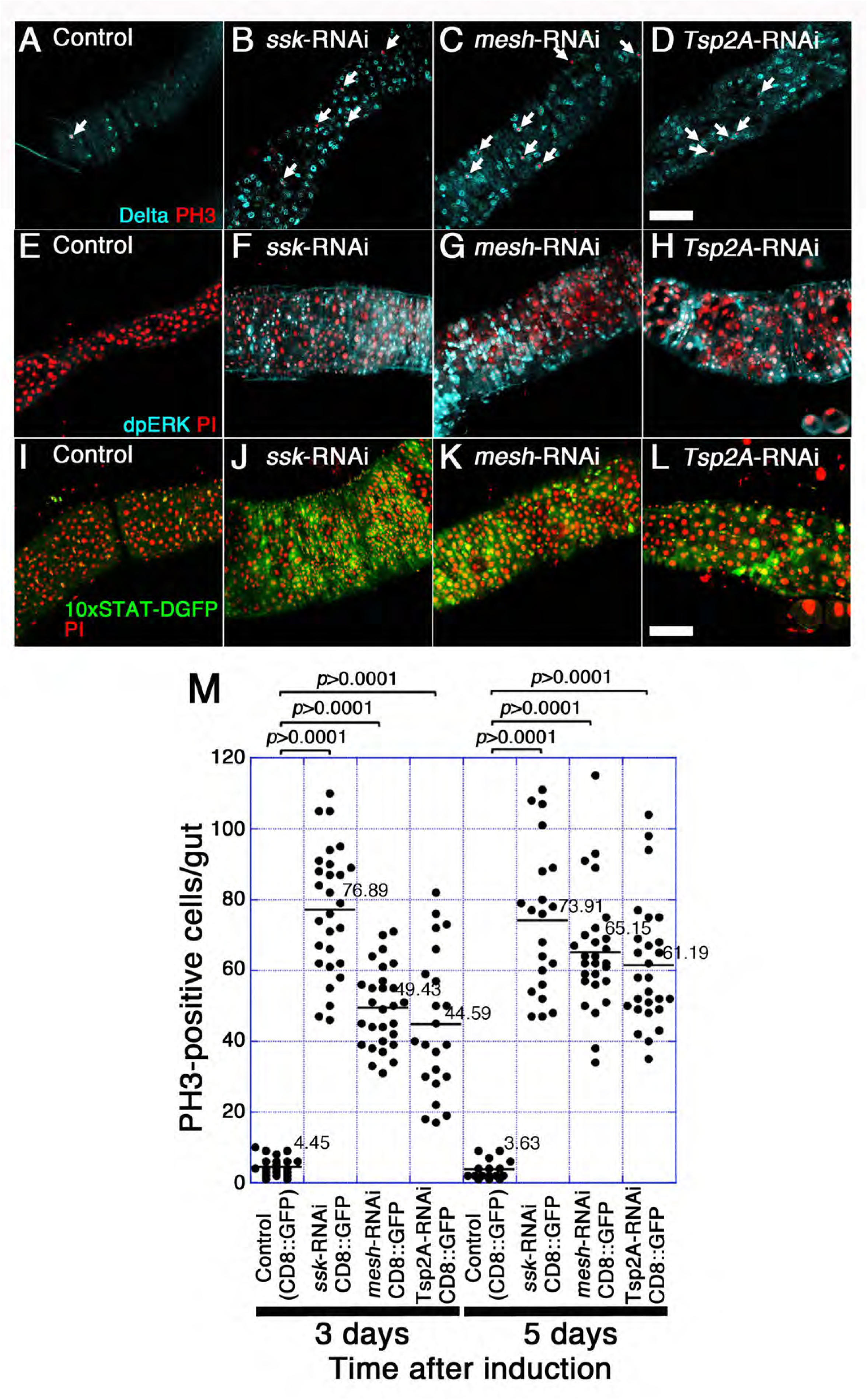
Depletion of sSJ-proteins from ECs leads to increased ISC proliferation in the midgut. **(A–H)** Confocal images of the adult anterior midgut expressing *Myo1A^ts^*-Gal4/UAS-*CD8-GFP* without (A, E, control) or with UAS-*ssk*-RNAi (B, F), UAS-*mesh*-RNAi (C, G), or UAS-*Tsp2A*-RNAi (D, H) at 5 days after induction stained for PH3 (red in A–D), Delta (blue in A–D), dpERK (blue in E–H), and DNA (propidium iodide) (red in E–H). PH3-positive cells and Delta-positive cells are increased in the sSJ-protein-deficient midgut compared with the control midgut (A–D). Enhancement of Ras-MAPK pathway activity in the sSJ-protein-deficient midgut is shown by increased expression of dpERK (E–H). Scale bar: 50 μm. **(I–L)** Confocal images of the adult anterior midgut expressing *Myo1A^ts^*-Gal4/*10xSTAT-DGFP* without (I, control) or with UAS-*ssk*-RNAi (J), UAS-*mesh*-RNAi (K), or UAS-*Tsp2A*-RNAi (L) at 5 days after induction stained for GFP (green) and DNA (propidium iodide) (red). Enhancement of JAK-STAT pathway activity in sSJ-protein deficient midgut is shown by increased expression of the *10xSTAT-DGFP* reporter. Scale bar: 50 μm. **(M)** Quantification of PH3-positive cells. The dot-plots show the numbers of PH3-positive cells in individual midguts. Left to right: Control (*CD8-GFP*) (*n*=22), *ssk*-RNAi *CD8-GFP* (*n*=28), *mesh*-RNAi *CD8-GFP* (*n*=28), and *Tsp2A*-RNAi *CD8-GFP* (*n*=27) at 3 days after induction, Control (*CD8-GFP*) (*n*=16), *ssk*-RNAi *CD8-GFP* (*n*=21), *mesh*-RNAi *CD8-GFP* (*n*=27), and *Tsp2A*-RNAi *CD8-GFP* (*n*=27) at 5 days after induction. Bars and numbers in the graph represent the mean PH3-positive cells in the fly lines. Statistical significance (*p*<0.0001) was evaluated by the Mann–Whitney *U*-test.

In the adult midgut, the Ras-MAP kinase and Jak-Stat signaling pathways are required for activation of ISC proliferation during the regeneration of epithelial cells (Buchon et al., 2010, Biteau and Jasper, 2011, Jiang and Edgar, 2009, Jiang et al., 2011, Osman et al., 2012, Zhou et al., 2013). Therefore, we examined whether these signaling pathways were activated in the sSJp-RNAis midgut. To monitor Ras-MAP kinase pathway activity, we examined the levels of phosphorylated ERK (dpERK) (Gabay et al., 1997). In control flies, dpERK signals were barely detectable in the midgut (Fig. 3E and Fig. S5A). In contrast, intense dpERK signals were observed in the sSJp-RNAis midgut, not only in cells residing on the basal side of the epithelium, but also in some ECs (Fig. 3F–H and Fig. S5B–D), strongly suggesting that the Ras-MAP kinase pathway was activated in ISCs, EBs, and certain ECs. To monitor Jak-Stat pathway activity, we used a Stat92E reporter line driving expression of DGFP (*10xSTAT-DGFP*). In the control midgut, few DGFP-positive cells were observed (Fig. 3I and Fig. S5E, I), while DGFP-positive cells were markedly increased on the basal side of the epithelium throughout the sSJp-RNAis midgut (Fig. 3J–L and Fig. S5F–H, J–L). In addition, dpERK- and DGFP-positive signals were detected in Esg-LacZ-positive cells in the *mesh*-RNAi midgut (Fig. S6), indicating that the MAP kinase and Jak-Stat pathways were activated in the progenitor cells in the sSJ-disrupted midgut. Collectively, these results demonstrate that depletion of sSJ-proteins from ECs results in activation of both the Ras-MAP kinase and Jak-Stat signaling pathways in the midgut.

### Simultaneous loss of *unpaired2* and *unpaired3* suppresses the abnormal accumulation of ECs in the *mesh*-deficient midgut

In ISC proliferation and EB differentiation, the Jak-Stat signaling pathway is activated by cytokines known as Unpaired (Upd) ligands (Jiang et al., 2009, Osman et al., 2012, Zhou et al., 2013). Upd2 and Upd3 were reported to contribute to increased ISC division in the midgut upon aging or exposure to stress (Osman et al., 2012). To examine whether Upd2 and Upd3 were involved in the increased ISC proliferation and abnormal accumulation of ECs observed in the sSJp-RNAis midgut, we suppressed *mesh* expression in the midgut of *upd2* and *upd3* double-mutant (*upd2,3^Δ^*) flies by expression of *mesh*-RNAi. In the *mesh*-RNAi *upd2,3^Δ^* midgut, a large number of mitotic cells were still observed and showed a similar level to the *mesh*-RNAi midgut (Fig. 4B, C, G), while only a few mitotic cells were detected in the control and *upd2,3^Δ^* midgut (Fig. 4A, G). Meanwhile, the expansion of the midgut observed in *mesh*-RNAi flies was significantly suppressed in *mesh*-RNAi *upd2,3^Δ^* flies. At 3 days after RNAi induction, the diameter of the most posterior region of the midgut in *mesh*-RNAi *upd2,3^Δ^* flies was significantly smaller than that in *mesh*-RNAi flies and was similar to the *upd2,3^Δ^* flies and control flies (Fig. 4H). At 5 days after RNAi induction, the suppressive effect of *upd2,3^Δ^* in *mesh*-RNAi flies in the midgut diameter appeared to be more remarkable, but it may include the influence of *upd2,3^Δ^* alone on the midgut because the diameter of the most posterior region of the midgut in *upd2,3^Δ^* flies is smaller than that in control flies in this condition (Fig. 4D–F, H). Accumulation of ECs was still observed in *mesh*-RNAi *upd2,3^Δ^* flies (Fig. 4F). The *upd2,3^Δ^* flies expressing *mesh*-RNAi in ECs exhibited a shortened lifespan, resembling that in *mesh*-RNAi flies (Fig. S7A). The midgut barrier dysfunction seen in *mesh*-RNAi flies was also observed in *mesh*-RNAi *upd2,3^Δ^* flies (Fig. S7B), suggesting that Upd2 and Upd3 do not contribute to the loss of barrier function in the midgut. Taken together, our observations suggest that induction of Upd2 and/or Upd3 expression is responsible for the aberrant behavior of ECs in the sSJp-RNAis midgut.

**Figure 4.**
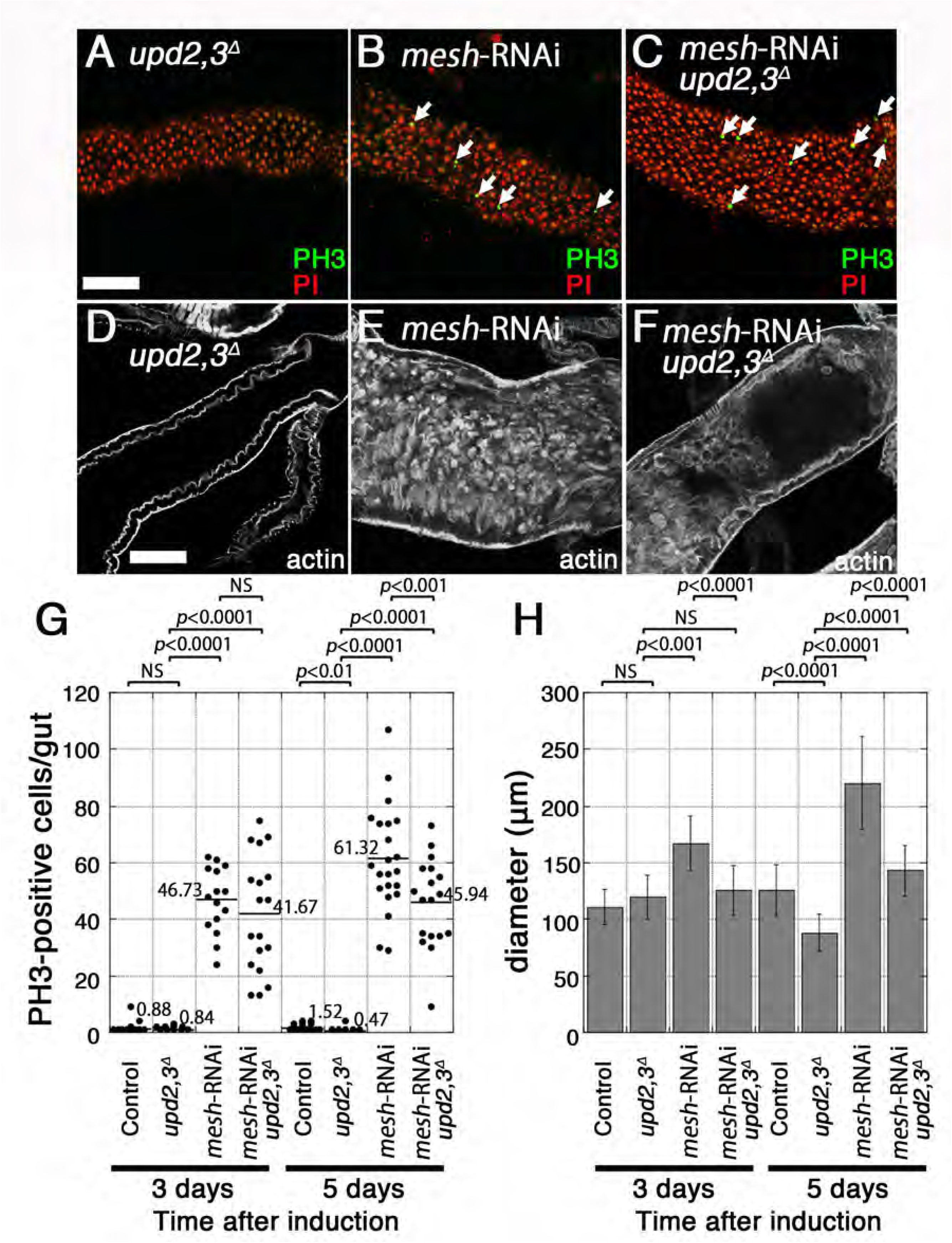
Loss of *upd2* and *upd3* suppresses abnormal accumulation of ECs in the *mesh*-deficient midgut. **(A–C)** Confocal images of the *upd2,3^Δ^* (A), *Myo1A^ts^*-Gal4/UAS-*mesh*-RNAi (B), and *Myo1A^ts^*-Gal4/UAS-*mesh*-RNAi *upd2,3^Δ^* (C) male fly anterior midgut at 5 days after induction stained for PH3 (green) and DNA (propidium iodide) (red). In the *mesh*-RNAi and *mesh*-RNAi *upd2,3^Δ^*midgut, PH3-positive cells are increased compared with the *upd2,3^Δ^* midgut. Scale bar: 50 μm. **(D–F)** Confocal images of the *upd2,3^Δ^*(D), *Myo1A^t^*^s^-Gal4/UAS-*mesh*-RNAi (E), and *Myo1A^ts^*-Gal4/UAS-*mesh*-RNAi *upd2,3^Δ^* (F) male fly midgut at 5 days after induction stained for actin (blue). The diameter of the most posterior region of the *mesh*-RNAi midgut is severely expanded compared with the *upd2,3^Δ^* midgut. The diameter of the *mesh*-RNAi *upd2,3^Δ^* posterior midgut is reduced compared with the *mesh*-RNAi midgut. Scale bar: 50 μm. **(G)** Quantification of PH3-positive cells in the *mesh*-RNAi and *mesh*-RNAi *upd2,3^Δ^* male fly midgut. The dot-plots show the numbers of PH3-positive cells in individual midguts. Left to right: Control (+/*mesh*-RNAi) (*n*=26), *upd2,3^Δ^* (*n*=19), *mesh*-RNAi (*Myo1A^ts^*/*mesh*-RNAi) (*n*=22), and *mesh*-RNAi *upd2,3^Δ^* (*upd2,3^Δ^, Myo1A^ts^* /*mesh*-RNAi) (*n*=18) at 3 and 5 days after induction. The bars and numbers in the graph represent the mean PH3-positive cells in the fly lines. Statistical significance was evaluated by the Mann–Whitney *U*-test. **(H)** Diameter of the most posterior region of Control (+/*mesh*-RNAi) (*n*=27), *upd2,3^Δ^* (*n*=16), *mesh*-RNAi (*Myo1A^ts^*/*mesh*-RNAi) (*n*=23), and *mesh*-RNAi *upd2,3^Δ^*(*upd2,3^Δ^, Myo1A^ts^*/*mesh*-RNAi) (*n*=19) male fly midgut at 3 and 5 days after induction. The diameter of the midgut was measured just anterior to the Malpighian tubules. Error bars show s.e.m. Statistical significance was evaluated by Student’s *t*-test.

### *Tsp2A*-mutant clones induce non-cell-autonomous stem cell proliferation

To further validate the effects of sSJ-protein depletion on the adult midgut, we generated mitotic clones that lacked *Tsp2A* and simultaneously expressed GFP in the ISC lineage using the mosaic analysis with a repressible cell marker (MARCM) system (Lee and Luo, 2001). The clone size, indicated by the number of GFP-positive cells per clone, of *Tsp2A*-mutant clones was comparable to that of control clones (Fig. 5A, B, G). However, an increased number of PH3-positive cells outside the mutant clones was observed in *Tsp2A*-mutant clones compared with control clones (Fig. 5A, B, H). These results indicate that loss of *Tsp2A* in ECs has a non-cell-autonomous effect on ISC proliferation. In addition, immunostaining with anti-Pdm1 and anti-Prospero (Pros) antibodies, which label ECs and EEs, respectively (Micchelli and Perrimon, 2006, Ohlstein and Spradling, 2006), revealed that *Tsp2A*-mutant clones contained differentiated ECs and EEs (Fig. 5C–F). These results suggest that loss of *Tsp2A* does not block differentiation of the ISC lineage.

**Figure 5.**
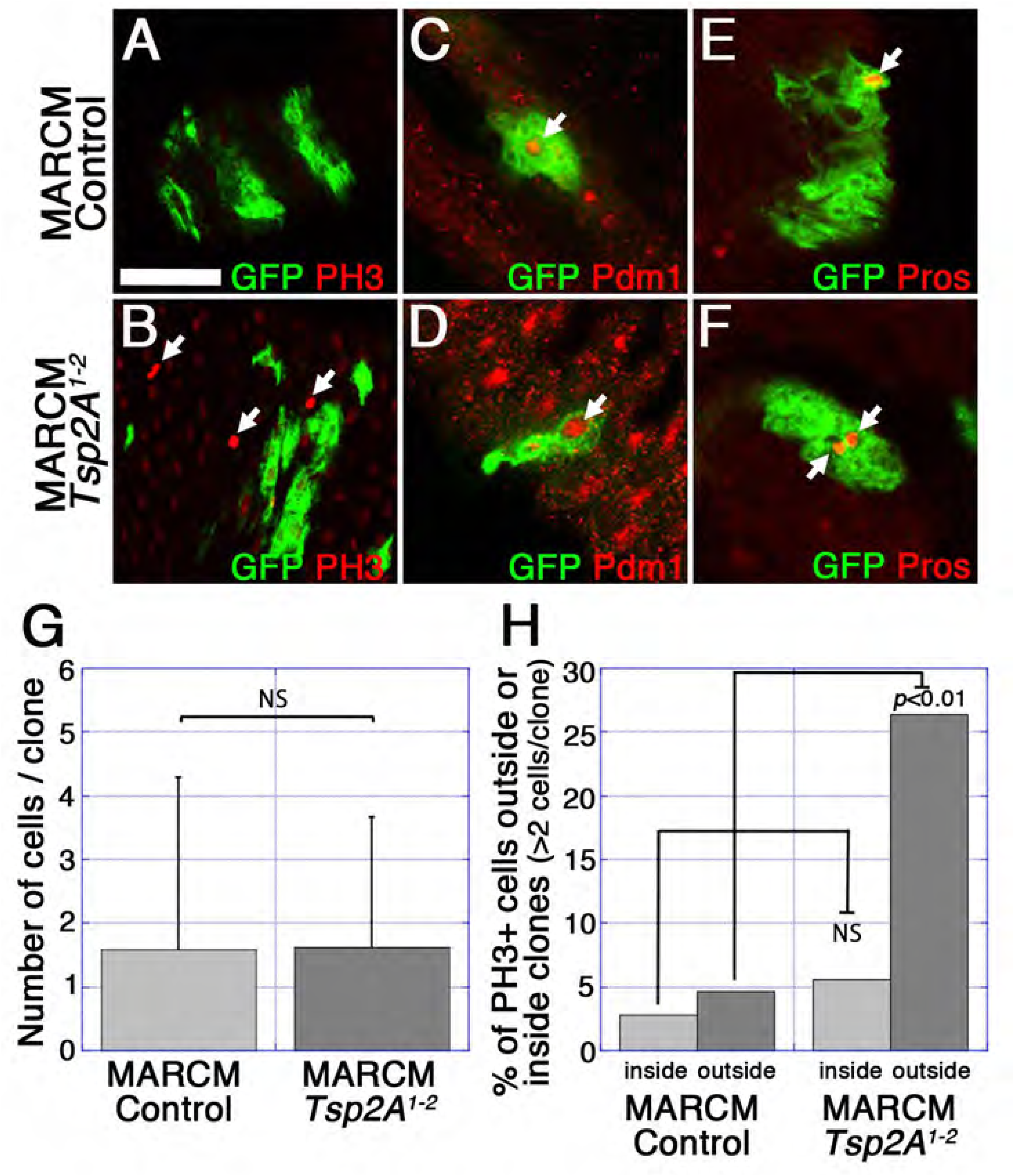
*Tsp2A*-mutant clones induce non-cell-autonomous stem cell proliferation. **(A–F)** Confocal images of the midgut from control clones (A, C, E) or *Tsp2A^1-2^* mutant clones (B, D, F) stained for GFP (green in A–F), PH3 (red in A, B), Pdm1 (red in C and D), and Pros (red in E and F). Control and *Tsp2A^1-2^* mutant clones were generated using the MARCM system and marked by GFP expression. The *Tsp2A^1-2^* mutant clones induce cell division of neighboring cells (arrows in B). In the *Tsp2A^1-2^*mutant clones, Pdm1- and Pros-positive cells are generated, similar to the control clones (arrows in C–F). Scale bar: 50 μm. **(G)** Quantification of average clone size for control and *Tsp2A^1-2^* mutant clones. Error bars show s.e.m. Statistical significance was evaluated by the Mann–Whitney *U*-test. NS, not significant. **(H)** Quantification of control and *Tsp2A^1-2^* mutant clones containing PH3-positive cells on the inside or outside of the clones. A total of 107 control clones and 197 *Tsp2A^1-2^* mutant clones were counted (each clone contained >2 cells). The *p*-value represents a significant difference in pairwise post-test comparisons indicated by the corresponding bars (Fisher’s exact test). NS, not significant.

## Discussion

In the *Drosophila* midgut epithelium, the paracellular barrier is constructed by specialized cell-cell junctions known as sSJs (Tepass and Hartenstein, 1994). Our previous studies revealed that three sSJ-associated membrane proteins, Ssk, Mesh, and Tsp2A, are essential for the organization and function of sSJs (Yanagihashi et al., 2012, Izumi et al., 2012, Izumi et al., 2016). In this study, we depleted the sSJ-proteins from ECs in the *Drosophila* adult midgut and showed that they are also required for the barrier function in the adult midgut epithelium. Interestingly, the reduced expression of sSJ-proteins in ECs led to remarkably shortened lifespan in adult flies, increased ISC proliferation, and intestinal hypertrophy accompanied by accumulation of morphologically aberrant ECs in the midgut. The intestinal hypertrophy caused by *mesh* depletion was suppressed by loss of *upd2* and *upd3* without profound suppression of ISC proliferation, recovery of the shortened lifespan, and recovery of the midgut barrier dysfunction. We also found that *Tsp2A*-mutant clones promoted ISC proliferation in a non-cell-autonomous manner. Taken together, we propose that sSJs play a crucial role in maintaining tissue homeostasis through regulation of ISC proliferation and EC behavior in the *Drosophila* adult midgut. The adult *Drosophila* intestine provides a powerful model to investigate the molecular mechanisms behind the emergence and progression intestinal metaplasia and dysplasia, which are associated with gastrointestinal carcinogenesis in mammals (Li and Jasper, 2016). Given that *Drosophila* intestinal dysplasia is associated with over-proliferation of ISCs and their abnormal differentiation, the intestinal hypertrophy observed in the present study should be categorized as a typical dysplasia in the *Drosophila* intestine.

Based on our observations, we hypothesize the following scenario for the hypertrophy generation in the sSJ-protein-deficient midgut. First, depletion of sSJ-proteins from ECs leads to disruption of sSJs in the midgut. Second, the impaired midgut barrier function caused by disruption of sSJs results in leakage of harmful substances from the intestinal lumen, thereby inducing the expression of cytokines and growth factors, such as Upd and EGF ligands, in the midgut. Alternatively, disruption of sSJs causes direct activation of a particular signaling pathway that induces expression of cytokines and growth factors by ECs. Third, proliferation of ISCs is promoted by activation of the Jak-Stat and Ras-MAP kinase pathways. Fourth, EBs produced by the asymmetric division of ISCs differentiate into ECs with impaired sSJs in response to cytokines such as Upd2 and/or Upd3. Fifth, the ECs fail to integrate into the epithelial layer because of insufficient cell-cell adhesion caused by impaired sSJs, and thus become stratified in the midgut lumen to generate hypertrophy. Interestingly, loss of *upd2* and *upd3* suppressed the intestinal hypertrophy caused by depletion of *mesh*, but not the increased ISC proliferation. These findings imply that Upd2 and/or Upd3 preferentially promote EB differentiation rather than ISC proliferation. Considering that Upd-Jak-Stat signaling is required for both ISC proliferation and EB differentiation (Jiang et al., 2009), Upd2 and/or Upd3 may predominantly promote EB differentiation and accumulation of ECs, while other cytokines such as Upd1 and/or EGF ligands may activate ISC proliferation in the sSJ-disrupted midgut. In this study, we observed abnormal morphology and aberrant F-actin and Dlg distributions in *ssk*-, *mesh*-, and *Tsp2A*-RNAi ECs. Consistent with our results, Chen et al. (2018) recently reported that loss of *mesh* and *Tsp2A* in clones causes defects in polarization and integration of ECs in the adult midgut. In contrast, no remarkable defects in the organization and polarity of ECs were observed in the *ssk*-, *mesh*-, and *Tsp2A*-mutant midgut in first-instar larvae (Yanagihashi et al., 2012, Izumi et al., 2012, Izumi et al., 2016), suggesting that sSJ-proteins are not required for establishment of the initial epithelial apical-basal polarity. This discrepancy may be explained by the marked difference between the larval and adult midguts: ECs in the larval midgut are postmitotic, while those in the adult midgut are capable of regeneration by the stem cell system (Lemaitre and Miguel-Aliaga, 2013). In the sSJ protein-deficient adult midgut, activated proliferation of ISCs generates excessive ECs. These ECs may lack sufficient cell-cell adhesion because of impaired sSJs, fail to become integrated into the epithelial layer, and detach from the basement membrane, leading to loss of normal polarity. Because sSJs seem to be the sole continuous intercellular contacts between adjacent epithelial cells in the midgut (Tepass and Hartenstein, 1994, Baumann, 2001), it is reasonable to speculate that sSJ-disrupted ECs have reduced cell-cell adhesion ability.

A recent study revealed that depletion of the tricellular junction protein Gliotactin from ECs leads to epithelial barrier dysfunction, increased ISC proliferation, and blockade of differentiation in the midgut of young adult flies (Resnik-Docampo et al., 2017). In contrast to the findings after depletion of sSJ-proteins in the present study, the *gliotactin*-deficient midgut does not appear to exhibit intestinal hypertrophy accompanied by accumulation of ECs throughout the midgut. Furthermore, the lifespan of *gliotactin*-deficient flies is longer than that of sSJ-protein-deficient flies. The difference in phenotypes between the two studies may reflect the difference in the degrees of sSJ deficiency: disruption of entire bicellular sSJs or tricellular sSJs only. Aging was also reported to be correlated with barrier dysfunction, increased ISC proliferation, and accumulation of aberrant cells in the adult midgut (Biteau et al., 2008, Rera et al., 2012). The hypertrophy formation in the sSJ-disrupted midgut accompanied by increased ISC proliferation and accumulation of aberrant ECs raise the possibility that disruption of sSJs is the primary cause of the alterations in the midgut epithelium with aging.

During the preparation of our manuscript, two groups published interesting phenotypes of the sSJ-protein-deficient adult midgut in *Drosophila* that are highly related to the present study. Salazar et al. (2018) reported that reduced expression of *ssk* in ECs leads to gut barrier dysfunction, altered gut morphology, increased stem cell proliferation, dysbiosis, and reduced lifespan. They also showed that up-regulation of Ssk in the midgut protects flies against microbial translocation, limits dysbiosis, and prolongs lifespan. Meanwhile, Xu et al. (2018) reported that depletion of *Tsp2A* from ISCs/EBs causes accumulation of ISCs/EBs and a swollen midgut with multilayered epithelium, similar to our observations. They also showed that knockdown of *ssk* and *mesh* in ISCs/EBs results in accumulation of ISCs/EBs. Importantly, they demonstrated that *Tsp2A* depletion from ISCs/EBs causes excessive aPKC-Yki-JAK-Stat activity and leads to increased stem cell proliferation in the midgut. They further showed that Tsp2A is involved in endocytic degradation of aPKC, which antagonizes the Hippo pathway. Their results strongly suggest that sSJs are directly involved in the regulation of intracellular signaling for ISC proliferation. In their study, *Tsp2A* knockdown in ISCs/EBs caused no defects in the midgut barrier function, in contrast to the present study. This discrepancy may be due to differences in the GAL4 drivers used in each study or the conditions for the barrier integrity assay/Smurf assay. In addition, Xu et al. (2018) mentioned that MARCM clones generated from ISCs expressing *Tsp2A*-RNAi grow much larger than control clones, while we found no remarkable size difference between Tsp2A mutant clones and control clones. Such discrepancies need to be reconciled by future investigations. To further clarify the mechanistic details for the role of sSJs in stem cell proliferation, it will be interesting to analyze the effects of sSJ-protein depletion on the behavior of adult Malpighian tubules, which also have sSJs, as well as on a stem cell system (Singh et al., 2007).

In this study, we have demonstrated that sSJs play a crucial role in maintaining tissue homeostasis through regulation of ISC proliferation and EC behavior in the *Drosophila* adult midgut. Our sequential identification of the sSJ-proteins Ssk, Mesh, and Tsp2A has provided a *Drosophila* model system that can be used to elucidate the roles of the intestinal barrier function by experimental dysfunction of sSJs in the midgut. However, as described in this study, simple depletion of sSJ-proteins throughout the adult midgut causes phenotypes that are too drastic, involving not only disruption of the intestinal barrier function but also intestinal dysplasia and subsequent lethality. To investigate the systemic effects of intestinal barrier impairment throughout the life course of *Drosophila*, more modest depletion of sSJ-proteins is needed for future studies.

## Material and Methods

### Fly stocks

Fly stocks were reared on a standard cornmeal fly medium at 25°C. *w^1118^* flies were used as wild-type flies unless otherwise specified. The other fly stocks used were: *y w*; *Myo1A*-GAL4 (#112001; Drosophila Genetic Resource Center (DGRC), Kyoto, Japan), *tubP*-GAL80*^ts^* (#7019; Bloomington Drosophila Stock Center (BDSC), Bloomington, IN), *y w*; CD8-GFP (#108068; DGRC), *y w*; *Pin^Yt^*/CyO; UAS-*mCD8-GFP* (#5130; BDSC), *w*; *10xStat92E-DGFP*/CyO (#26199; BDSC), *w*; *10xStat92E-DGFP*/TM6C *Sb Tb* (#26200; BDSC), *y w*; *esg-lacZ*/CyO (#108851; DGRC), FRT19A; *ry* (#106464; DGRC).

The RNAi lines used were: *ssk*-RNAi (Yanagihashi et al., 2012), *mesh*-RNAi (#12074R-1, 12074R-2; Fly Stocks of National Institute of Genetics (NIG-Fly), Mishima, Japan) (Izumi et al., 2012), *Tsp2A*-RNAi (#11415R-2; NIG-Fly *Tsp2A*IR1-2) (Izumi et al., 2016), *w*-RNAi (#28980; BDSC).

The mutant stocks used were: *Tsp2A^1-2^* (Izumi et al., 2016), *w upd2^Delta^ upd3^Delta^* (#55729; BDSC).

The following stocks were used to generate positively-marked MARCM clones: *tub^P^*-GAL80 *w* FRT19A; *Act5C*-GAL4, UAS-*GFP*/CyO (#42726; BDSC), FRT19A *tub^P^*-GAL80 hsFLP *w*; UAS-*mCD8GFP* (#108065; DGRC).

### Conditional expression of UAS transgenes (TARGET system)

Flies are crossed and grown at 18°C until eclosion. Adult flies at 2–5 days after eclosion were collected and transferred to 29°C for inactivation of Gal80. All analyses for these experiments were performed on female flies, because their age-related gut pathology is well established (Lemaitre and Miguel-Aliaga, 2013).

### MARCM clone induction

Flies were crossed at 18°C. At 2–5 days after eclosion, adult flies were heat-shocked at 37°C for 1 h twice daily. Adult flies were transferred to fresh vials every 2–3 days and maintained at 25°C for 10 days after clone induction before being dissected.

### Immunostaining

Adult flies were dissected in Hanks’ Balanced Salt Solution and the midgut was fixed with 4% paraformaldehyde in PBS/0.2% Tween-20 for 30 min. The fixed specimens were washed three times with PBS/0.4% Triton X-100 and blocked with 5% skim milk in PBS/0.2% Tween-20. Thereafter, the samples were incubated with primary antibodies at 4°C overnight, washed three times with PBS/0.2% Tween-20, and incubated with secondary antibodies for 3 h. After another three washes, the samples were mounted in Fluoro-KEEPER (12593-64; Nakalai Tesque, Kyoto, Japan). Images were acquired with a confocal microscope (Model TCSSPE; Leica Microsystems, Wetzlar, Germany) using its accompanying software and HC PLAN Apochromat 20× NA 0.7 and HCX PL objective lenses (Leica Microsystems). Images were processed with Adobe Photoshop® software (Adobe Systems Incorporated, San Jose, CA).

### Antibodies

The following primary antibodies were used: rabbit anti-Mesh (955-1; 1:1000) (Izumi et al., 2012); rabbit anti-Tsp2A (302AP; 1:200) (Izumi et al., 2016); rabbit anti-Ssk (6981-1; 1:1000) (Yanagihashi et al., 2012); mouse anti-Dlg (4F3; Developmental Studies Hybridoma Bank (DSHB), Iowa City, IA; 1:50); mouse anti-Delta (C594.9B; DSHB; 1:20); mouse anti-Pros (MR1A; DSHB; 1:20); rabbit anti-Pdm1 (a gift from Dr. Yu Cai; 1:500) (Yeo et al., 1995); rabbit anti-PH3 (06-570; Millipore, Darmstadt, Germany; 1:1000); rabbit anti-dpERK (Cell Signaling, Danvers, MA; 1:500); rat anti-GFP (GF090R; Nakalai Tesque; 1:1000); rabbit anti-GFP (598; MBL, Nagoya, Japan; 1:1000), and mouse anti-β-galactosidase (Z3781; Promega, Madison, WI; 1:200). Alexa Fluor 488-conjugated (A21206; Thermo Fisher, Waltham, MA) and Cy3- and Cy5-conjugated (Jackson ImmunoResearch Laboratories, West Grove, PA) secondary antibodies were used at 1:400 dilution. Actin was stained with Alexa Fluor 568 phalloidin (A12380; Thermo Fisher; 1:1000) or Alexa Fluor 647 phalloidin (A22287; Thermo Fisher; 1:1000). Nuclei were stained with propidium iodide (Nakalai Tesque; 0.1 mg ml^-1^).

### Electron microscopy and toluidine blue staining

Adult control, *ssk*-, *mesh-*, and *Tsp2A-*RNAi flies at 5 days after transgene induction were dissected and fixed overnight at 4°C with a mixture of 2.5% glutaraldehyde and 2% paraformaldehyde in 0.1 M cacodylate buffer (pH 7.4). The specimens including the midgut were prepared as described previously (Izumi et al., 2012). Ultrathin sections (50–100 nm) were double-stained with 4% hafnium (IV) chloride and lead citrate, and observed with a JEM-1011 electron microscope (JEOL, Tokyo, Japan) at an accelerating voltage of 80 kV. For toluidine blue staining, sections (0.5–1 μm) mounted on glass slides were placed in 0.05% toluidine blue in distilled water for 2–30 min, rinsed in water for 1 min, and allowed to air dry. The stained sections were observed with an optical microscope (BX41; Olympus, Tokyo, Japan).

### Statistical analyses

Statistical significance was evaluated by the Mann–Whitney *U*-test, Student’s *t*-test (KaleidaGraph software; Synergy Software, Reading, PA), and Fisher’s exact test. Values of *p*<0.05 were considered significant.

### Barrier integrity assay (Smurf assay)

Flies at 2–5 days of age were placed in empty vials containing a piece of paper soaked in 1% (wt/vol) Blue Dye No. 1 (Tokyo Chemical Industry, Tokyo, Japan)/5% sucrose solution at 50–60 flies/vial. After 2 days at 18°C, the flies were placed in new vials containing paper soaked in BlueDye/sucrose and transferred to 29°C. Loss of midgut barrier function was determined when dye was observed outside the gut (Rera et al., 2011, Rera et al., 2012). Flies were transferred to new vials every 2 days.

## Acknowledgments

We are grateful to all members of the Furuse laboratories for helpful discussions. We thank the Bloomington Drosophila Stock Center, Drosophila Genetic Resource Center at Kyoto Institute of Technology, and Fly Stocks of National Institute of Genetics (NIG-Fly) for fly stocks. We also thank Alison Sherwin, PhD, from Edanz Group (www.edanzediting.com/ac) for editing a draft of this manuscript.

## Author contributions

Y.I. designed the research; Y.I. and K.F. performed the experiments; Y.I. analyzed the data; Y.I. and M.F. wrote the paper.

---

**Figure S1.**
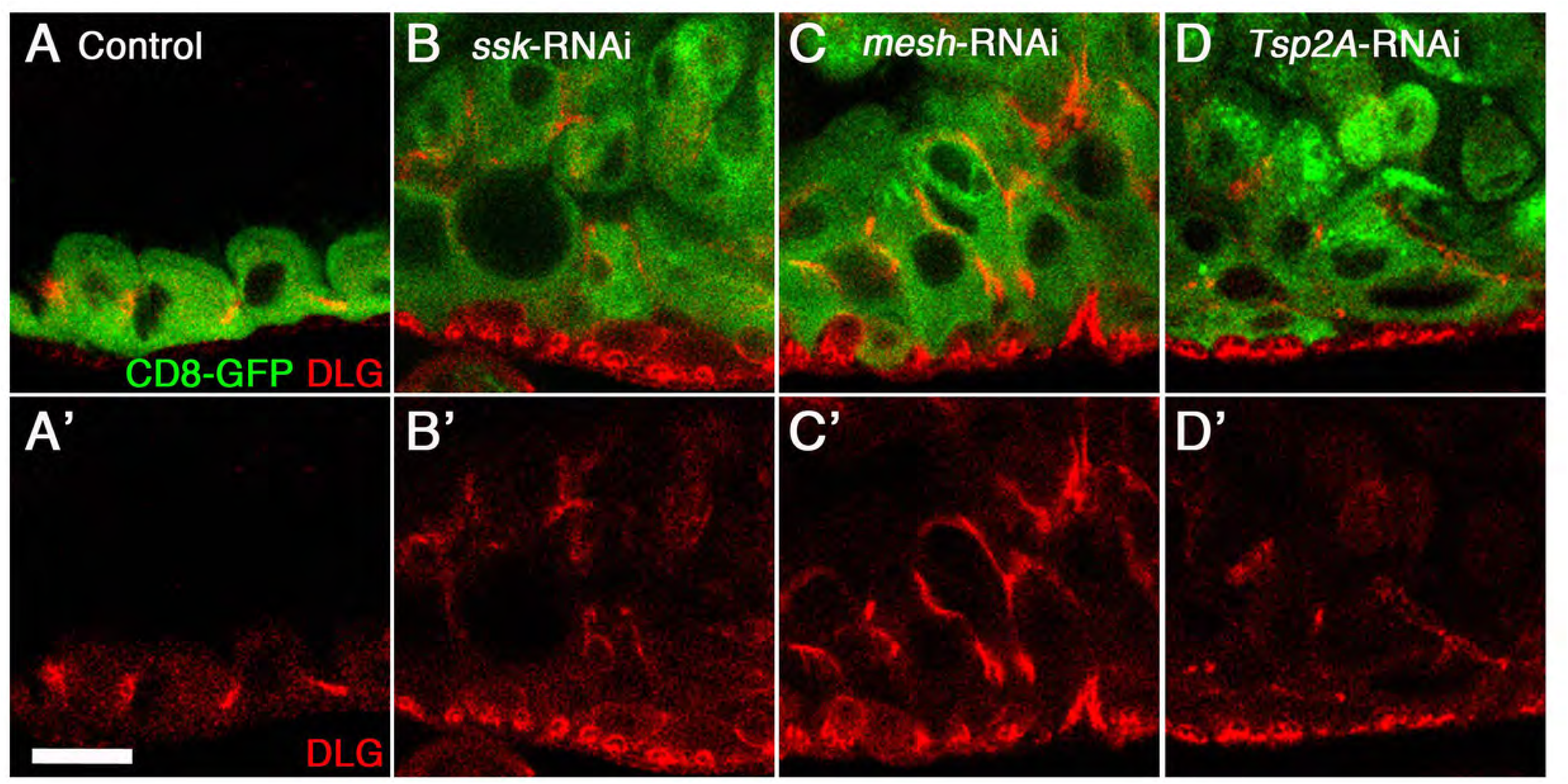
Dlg is mislocalized in sSJ-protein-deficient ECs. **(A–D’)** Confocal images of the adult anterior midgut expressing *Myo1A^ts^*-Gal4/UAS-*CD8-GFP* without (A, A’, control) or with UAS-*ssk*-RNAi (B, B’), UAS-*mesh*-RNAi (C, C’), or UAS-*Tsp2A*-RNAi (D, D’) at 5 days after induction stained for Dlg (red). The images show an optical cross-section through the center of the midgut. Scale bar: 20 μm.

**Figure S2.**
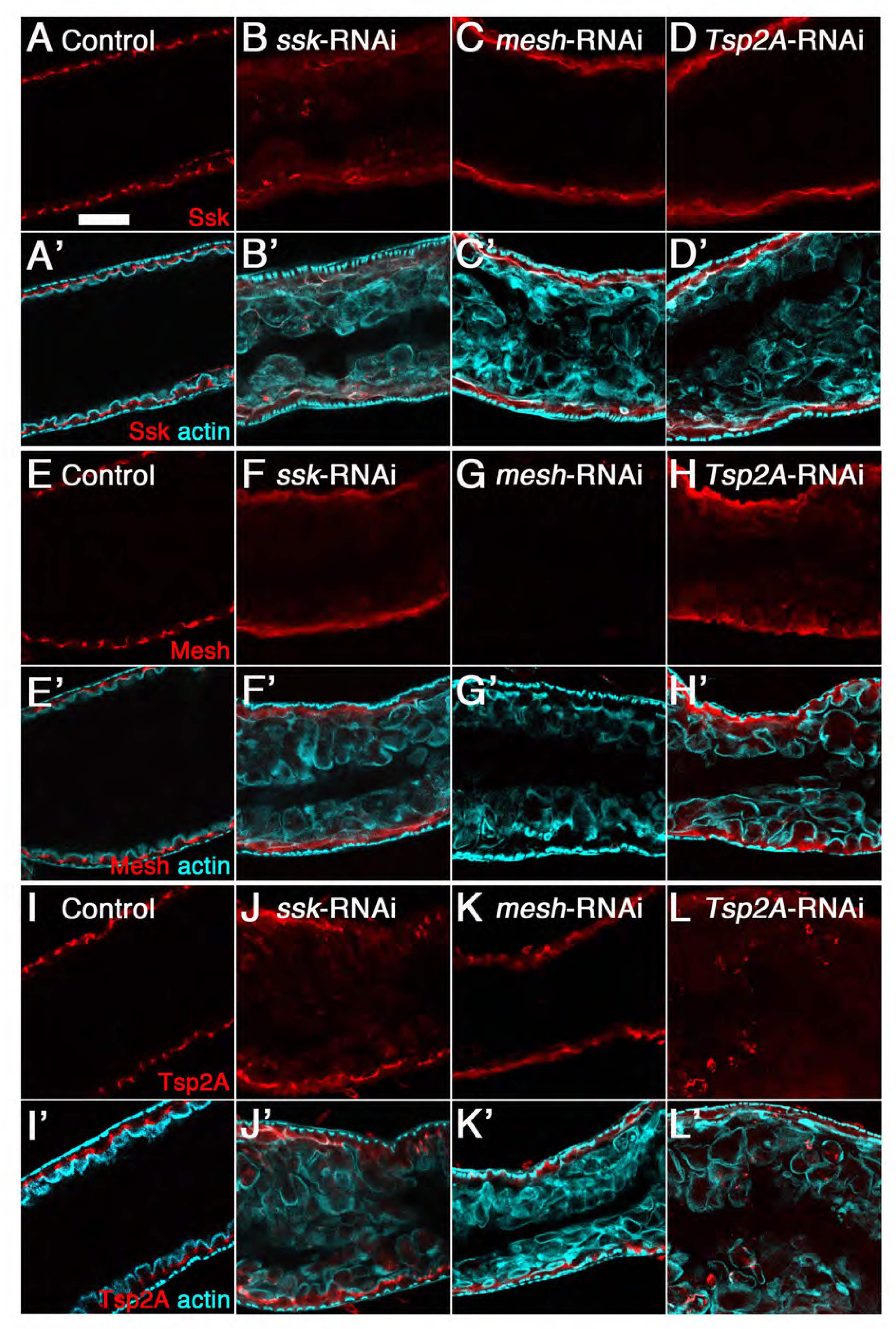
Expression of sSJp-RNAis in ECs leads to decreased levels of the respective target proteins and mislocalization of other sSJ-proteins in the midgut. **(A–L’)** Confocal images of the adult anterior midgut expressing *Myo1A^ts^*-Gal4/UAS-*CD8-GFP* without (A, A’, E, E’, I, I’, control) or with UAS-*ssk*-RNAi (B, B’, F, F’, J, J’), UAS-*mesh*-RNAi (C, C’, G, G’ K, K’), or UAS-*Tsp2A*-RNAi (D, D’, H, H’, L, L’) at 5 days after induction stained for Ssk (red in A–D), Mesh (red in E–H), Tsp2A (red in I–L), and actin (blue in A’–L’). Scale bar: 50 μm.

**Figure S3.**
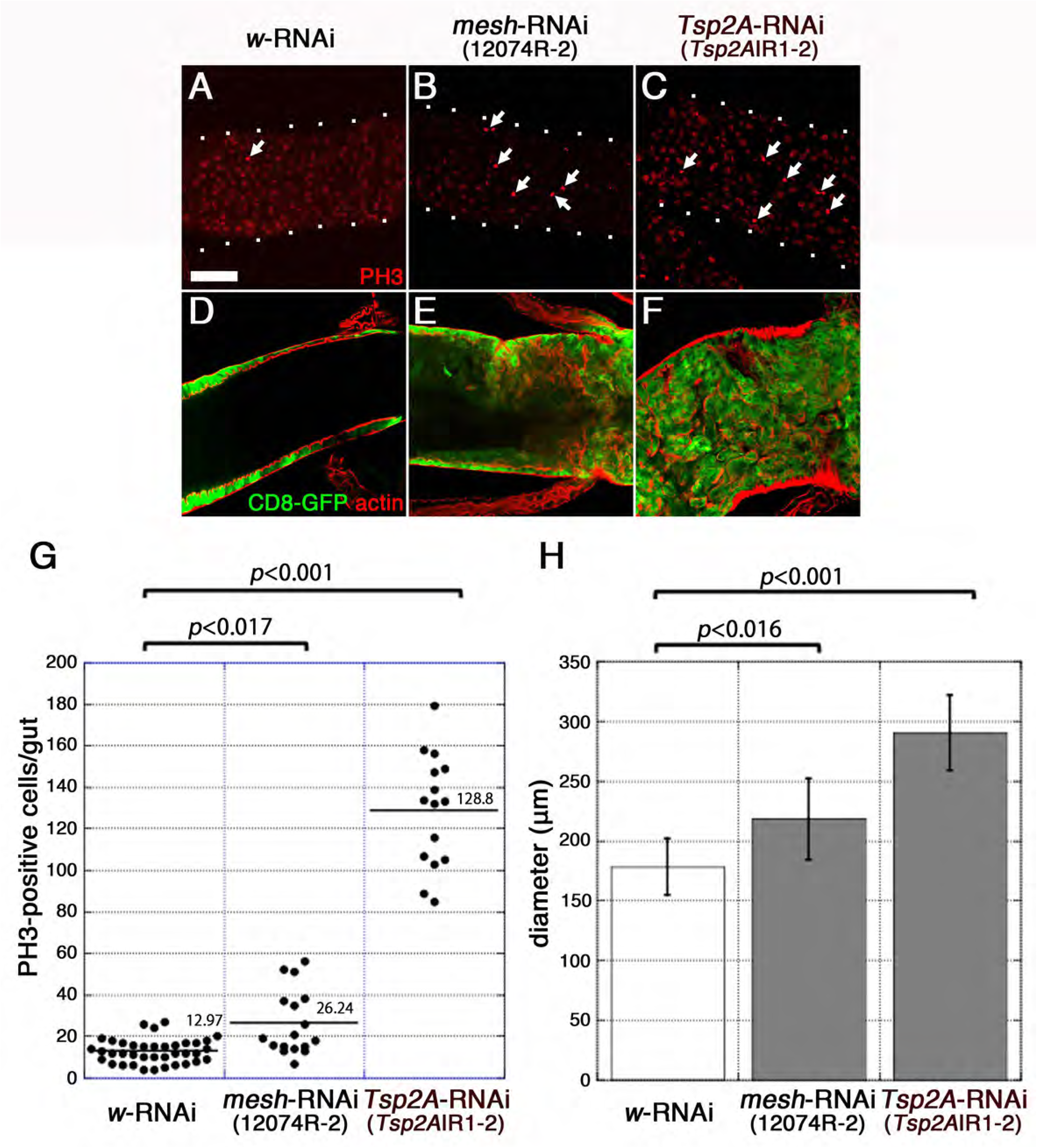
Expression of additional RNAi lines of *mesh* and *Tsp2A* in the adult midgut causes increased ISC proliferation and intestinal hypertrophy. **(A–F)** Confocal images of the adult anterior (A–C) and posterior (D–F) midgut expressing *Myo1A^ts^*-Gal4/UAS-*CD8-GFP* with UAS-*white*-RNAi (JF01545) (A, D, control), UAS-*mesh*-RNAi (12074R-2) (B, E), or UAS-*Tsp2A*-RNAi (*Tsp2A*IR1-2) (C, F) at 5 days after induction stained for PH3 (red in A–C) and actin (red in D–F). The dots in (A–C) show the outline of the midgut. Scale bar: 50 μm. **(G)** Quantification of PH3-positive cells. The number of PH3-positive cells per midgut was counted at 5 days after induction. The dot-plots show the numbers of PH3-positive cells in individual midguts. Left to right: Control (*white*-RNAi *CD8-GFP*) (*n*=38), *mesh*-RNAi *CD8-GFP* (*n*=17), and *Tsp2A*-RNAi *CD8-GFP* (*n*=15). The bars and numbers in the graph represent the mean PH3-positive cells in the fly lines. Statistical significance was evaluated by the Mann–Whitney *U*-test. **(H)** Diameter of the most posterior region of the midgut. The diameter of the midgut was measured just anterior to the Malpighian tubules at 5 days after induction. Left to right: Control (*white*-RNAi *CD8-GFP*) (*n*=23), *mesh*-RNAi *CD8-GFP* (*n*=14), and *Tsp2A*-RNAi *CD8-GFP* (*n*=17). Error bars show s.e.m. Statistical significance was evaluated by Student’s *t*-test.

**Figure S4.**
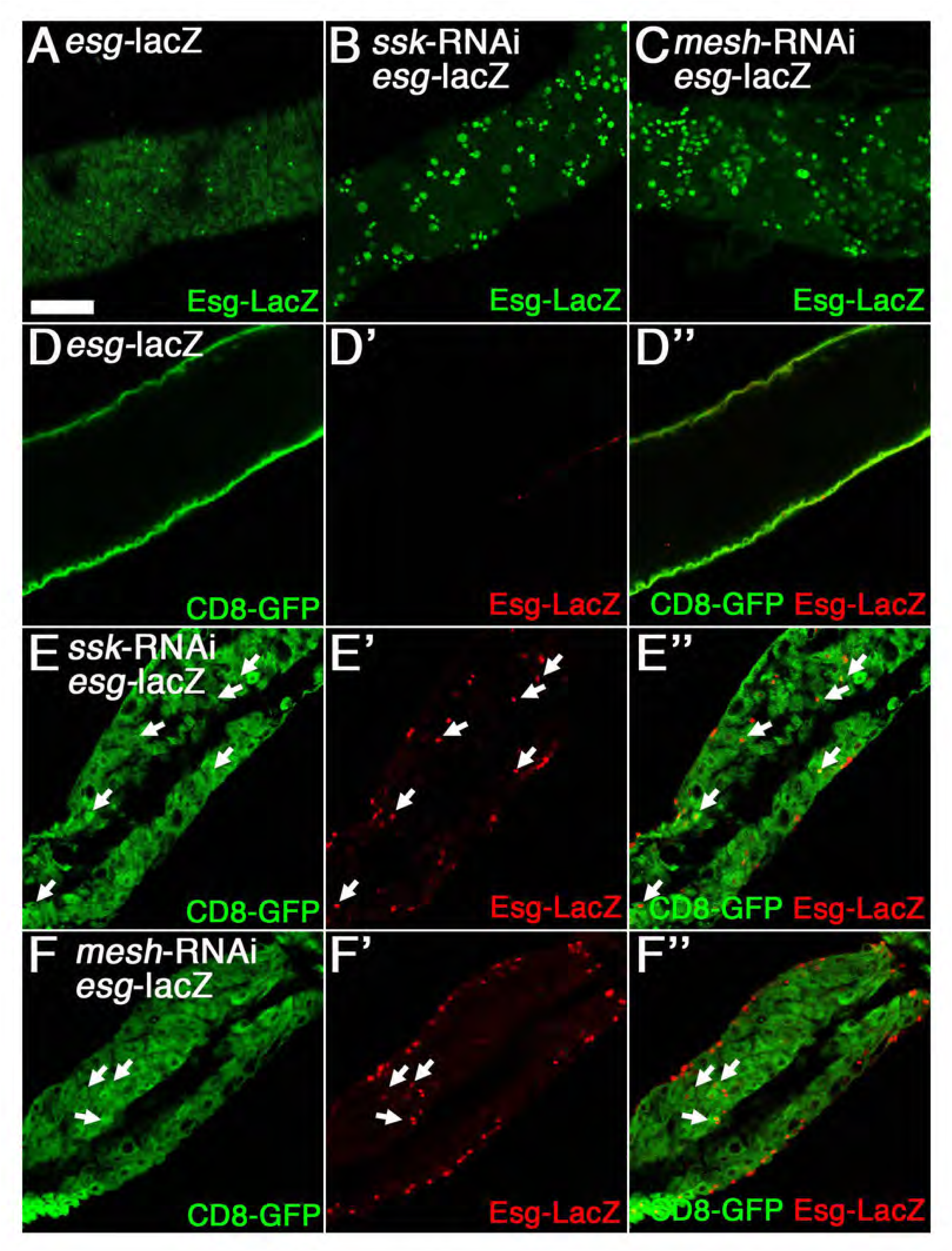
Depletion of *ssk* and *mesh* from ECs leads to an increase in Esg-LacZ-positive cells. **(A–F’’)** Confocal images of the adult anterior midgut expressing *Myo1A^ts^*-Gal4/UAS-*CD8-GFP/esg-lacZ* without (A, D, D’, D’’, control) or with UAS-*ssk*-RNAi (B, E, E’, E’’) or UAS-*mesh*-RNAi (C, F, F’, F’’) at 5 days after induction stained for β-galactosidase (green in A–C, red in D’, D’’, E’, E’’, F’, F’’). The images show an optical cross-section through the center of the midgut (D–F’’). The arrows in D–F’’ indicate CD8-GFP-driven *Myo1A*-GAL*4* and Esg-LacZ double-positive cells. Scale bar: 50 μm.

**Figure S5.**
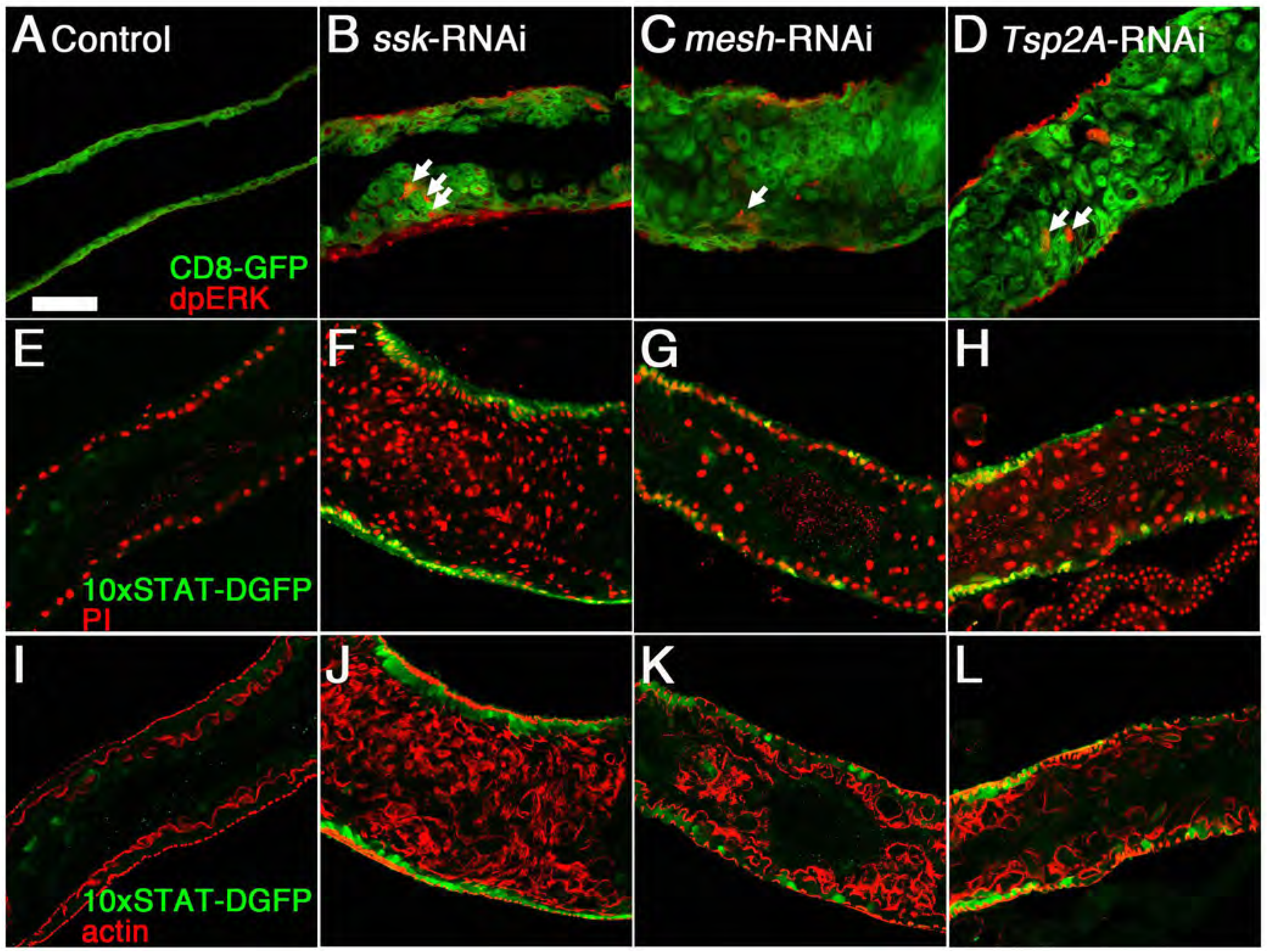
The Ras-MAP kinase pathway is activated in some ECs in the sSJ-protein-deficient midgut. **(A–D)** Confocal images of the adult anterior midgut expressing *Myo1A^ts^*-Gal4/UAS-*CD8-GFP* without (A, control) or with UAS-*ssk*-RNAi (B), UAS-*mesh*-RNAi (C), or UAS-*Tsp2A*-RNAi (D) at 5 days after induction stained for dpERK (red). The arrows indicate CD8-GFP-driven *Myo1A*-GAL4 (green) and dpERK double-positive cells. Scale bar: 50 μm. **(E–L)** Confocal images of the adult anterior midgut expressing *Myo1A^ts^*-Gal4/*10xSTAT-DGFP* without (E, I, control) or with UAS-*ssk*-RNAi (F, J), UAS-*mesh*-RNAi (G, K), or UAS-*Tsp2A*-RNAi (H, L) at 5 days after induction stained for GFP (green in E–L), DNA (propidium iodide) (red in E–H), and actin (red in I–L). (E and I), (F and J), (K and L), and (H and L) are each derived from the same sample. The images show an optical cross-section through the center of the midgut (A–L). Scale bar: 50 μm.

**Figure S6.**
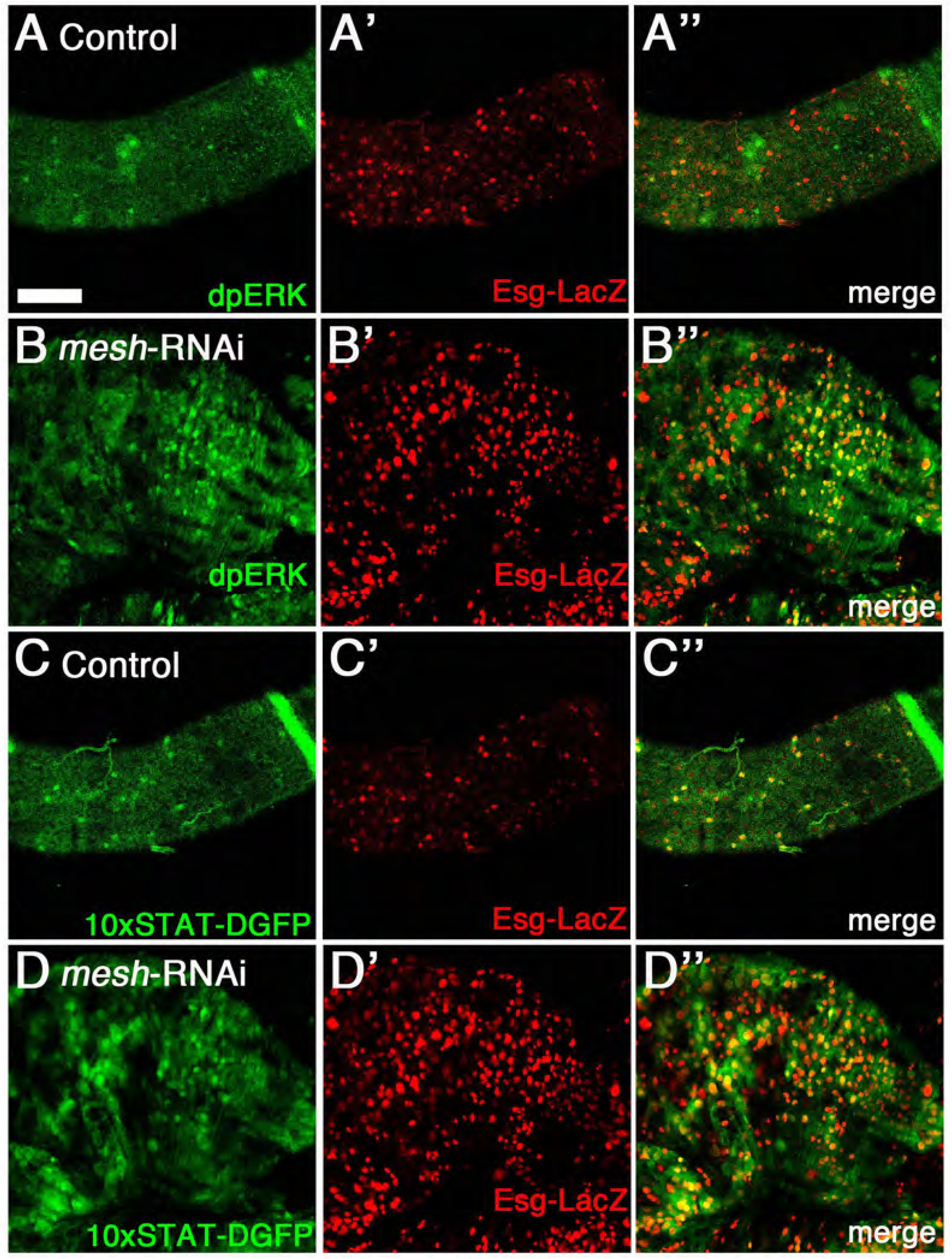
The Ras-MAP kinase and Jak-Stat pathways are activated in Esg-LacZ-positive cells in the *mesh*-deficient midgut. **(A–D’’)** Confocal images of the adult posterior midgut expressing *Myo1A^ts^*-Gal4/UAS-*CD8-GFP*/*esg-lacZ* without (A–A’’, C–C’’, control) or with UAS-*mesh*-RNAi (B–B’’, D–D’’) at 5 days after induction stained for β-galactosidase (red in A’, A’’, B’, B’’, C’, C’’, D’, D’’), dpERK (green in A, A’’, B, B’’), and GFP (green in C, C’’, D, D’’). (A and C) and (B and D) are each derived from the same sample. Scale bar: 50 μm.

**Figure S7.**
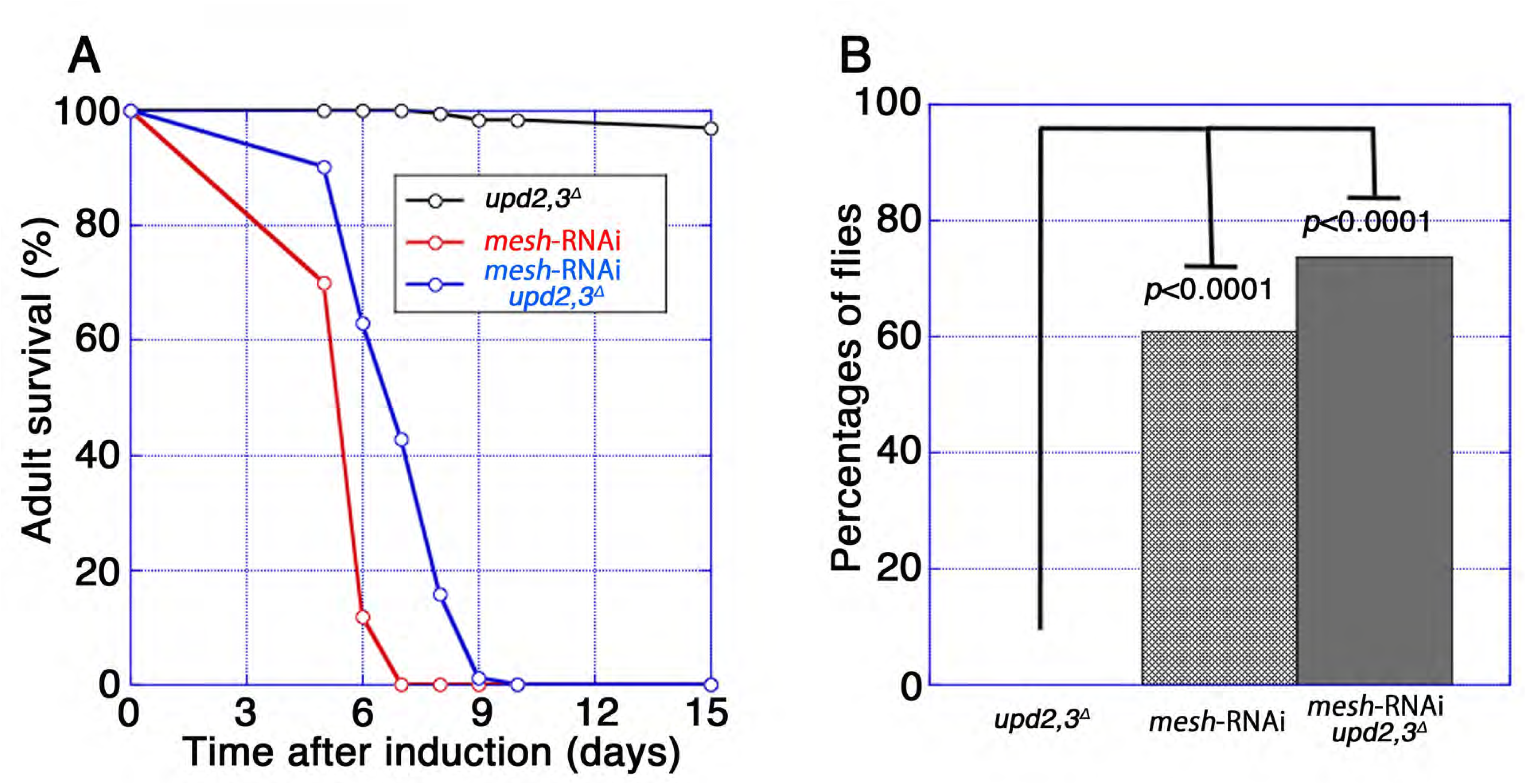
Depletion of *mesh* from ECs results in shortened lifespan and midgut barrier dysfunction in *upd2,3^Δ^* mutant flies. **(A)** Survival analysis of *upd2,3^Δ^*(*n*=240), *Myo1A^ts^*-Gal4/UAS-*mesh*-RNAi (*n*=180), and *Myo1A^ts^*-Gal4/UAS-*mesh*-RNAi *upd2,3^Δ^* (*n*=260) male flies. Each vial contained 20 male flies. **(B)** Barrier integrity assays (Smurf assays). In these assays, *upd2,3^Δ^* (*n*=209), *Myo1A^ts^*-Gal4/UAS-*mesh*-RNAi (*n*=192), and *Myo1A^ts^*-Gal4/UAS-*mesh*-RNAi *upd2,3^Δ^* (*n*=152) male flies were fed blue dye in sucrose solution. At 5 days after induction, the phenotypes were examined. The *p*-values represent significant differences in pairwise post-test comparisons indicated by the corresponding bars (Fisher’s exact test).

